# Golden Promise-rapid, a fast-cycling barley genotype with high transformation efficiency

**DOI:** 10.1101/2025.10.31.685778

**Authors:** Gabriele Buchmann, Einar Baldvin Haraldsson, Rebekka Schüller, Thea Rütjes, Agatha Alexandra Walla, Maria von Korff Schmising, Shanda Liu

## Abstract

The spring barley cultivar Golden Promise (GP) is the major reference genotype for transformation due to its high transformability and availability of a reference genome. However, GP is characterized by a long generation cycle and stress susceptibility under non-optimal growth conditions because it carries a mutation at the floral inducer *Photoperiod-H1* (*Ppd-H1*). Previously, we showed that a GP introgression line, Golden Promise-fast (GP-fast), generated by introducing the wild-type *Ppd-H1* allele from the winter barley cultivar Igri, exhibits early flowering and improved stress resilience. In this study, we generated a fast-cycling genotype, Golden Promise-rapid (GP-rapid), isogenic to GP with high transformation efficiency. We conducted two backcrosses of GP-fast to reduce the residual Igri genome. The resulting genotype contains only a single introgression of approximately 0.6 Mbp at the *Ppd-H1* locus on chromosome 2H. Under speed breeding conditions, its generation time was reduced to 63 days (25% shorter than GP’s 84 days). Parallel transformation of GP, GP-fast, and GP-rapid using CRISPR/Cas9-mediated genome editing of *Ppd-H1* revealed high regeneration and transformation efficiencies of GP-rapid, comparable to GP. Overall, we report on the development of a fast-cycling GP isogenic line as a research tool for efficient generation of transgenic and gene-edited barley plants.

**Highlights:** A new fast-cycling barley genotype, GP-rapid, reduces generation time by 25% while retaining high transformation efficiency, advancing functional genomic studies in barley.

## Introduction

Barley (*Hordeum vulgare*) is one of the major cereal crops and has emerged as a key genetic model for the Triticeae tribe. Its diploid genome, abundance of genetic resources, availability of a pan-genome, and transformation amenability make it particularly valuable for functional genomics (Saisho and Takeda, 2011; Harwood, 2019; Jayakodi et al., 2020). Recent advances in genome editing technologies have enabled the introduction of precise genetic modifications in barley (Han et al., 2021; Lawrenson et al., 2021; Tamilselvan-Nattar-Amutha et al., 2023; Lawrenson et al., 2024), providing a powerful tool to dissect causal relationships between genotype and phenotype.

Golden Promise (GP), a gamma-ray-induced mutant of the cultivar Maythorpe, has served as the reference genotype for barley transformation for over two decades because it exhibits high *Agrobacterium*-mediated transformation efficiency and strong regeneration capacity in tissue culture (Forster, 2001; Murray et al., 2004; Hensel et al., 2008; Harwood, 2012). Transformation attempts using other barley varieties have either failed or delivered low transformation efficiencies (Hensel et al., 2008; Harwood, 2012; Marthe et al., 2015). The use of GP for transformation and genome editing is further supported by the recent publication of a full reference genome (Schreiber et al., 2020; Jayakodi et al., 2020).

Despite these advantages, GP is also characterized by high environmental plasticity and a long generation cycle. GP carries a loss-of-function mutation in HvDep1, the Gγ subunit of a heterotrimeric G protein that positively regulates culm elongation and seed size in barley (Wendt et al., 2016). The authors demonstrated that the HvDep1 loss-of-function mutation conferred either an increase or a decrease in harvestable yield depending on the environment and genetic background, and thus enhances environmental plasticity. Previous studies reported that using stress-free explants is critical for achieving high transformation efficiency in GP (Harwood, 2014; Hayta et al., 2021). Its environmental plasticity poses challenges when donor plants are grown in non-optimal conditions. Additionally, as a spring barley, GP does not require vernalization to flower due to a deletion in the first intron of the vernalization gene *Vrn-H1* (5H) and a deletion of all three ZCCT copies underlying the *Vrn-H2* locus on 4H (Cockram et al., 2007). GP also carries a natural mutation at the major photoperiod response gene *Ppd-H1*, which causes a delay in reproductive development under long-day (LD) conditions (Turner et al., 2005). When grown under controlled conditions with 16h light, GP requires approximately 60-70 days to flower and 4 months to reach maturity (Campoli et al., 2012; Mulki et al., 2018; Pieper et al., 2021), which limits the number of generations to three per year. GP’s long generation cycle limits its suitability for the rapid generation of transgenic and gene-edited plants.

We have previously shown that introgressing the wild-type *Ppd-H1* allele from the winter barley cultivar Igri into GP accelerated flowering time by two weeks, which considerably shortened the generation time (Gol et al., 2021; Helmsorig et al., 2024). Furthermore, the resulting introgression line, termed GP-fast, displayed improved culm elongation and high developmental stability and spike fertility in response to heat and drought, in contrast to GP (Gol et al., 2021; Lan et al., 2025). GP-fast has been successfully used for generating CRISPR/Cas9 mutant lines (Amanda et al., 2022; Müller et al., 2023; Wanke et al., 2023; Helmsorig et al., 2024; Lan et al., 2025; Vardanega et al., 2025).

However, GP-fast retains at least three major Igri-derived genomic introgressions: one flanking the *Ppd-H1* locus on chromosome (Chr) 2H, and two others on Chr 6H and 7H (Gol et al., 2021). This might become problematic for designing genome editing guides because nucleotide differences between the introgression line and the GP reference sequence can reduce guide accuracy and may also affect the performance of resulting transgenic lines. Furthermore, the introgressions on 2H and 6H partially overlap with minor QTLs affecting the transformation amenability in GP (Hisano and Sato, 2016; Hisano et al., 2017).

In this work, we greatly reduced the donor genome fractions in GP-fast by performing two additional rounds of backcrosses to GP. Genotyping results showed that the newly generated genotype, GP-rapid, contains only a 0.6 Mbp Igri introgression surrounding the *Ppd-H1* locus on Chr 2H with only 26 genes within this region. GP-rapid completes one generation in 63 days under speed breeding conditions – a 25% reduction compared to GP. Importantly, GP-rapid showed comparable regeneration and transformation efficiencies to GP under our experimental conditions.

## Materials and methods

### Plant materials

Barley (*Hordeum vulgare*) genotypes used in this study included Golden Promise (GP), one selected Golden Promise-fast line (GP-fast_9, a sister line to the genotype described in Gol et al., 2021), and the newly developed line, Golden Promise-rapid (GP-rapid). GP-fast was generated by crossing GP with the winter barley cultivar Igri followed by three cycles of backcrossing and subsequent selfing (BC3F5) to obtain a homozygous line carrying the wild-type *Ppd-H1* allele. Chromosome-scale sequence assemblies of Golden Promise and Igri have been generated and compared within the barley pangenome project (Jayakodi et al., 2020).

GP, Igri, and the selected GP-fast line were first genotyped using the Barley 50k iSelect SNP Array (TraitGenetics GmbH, Germany) (Bayer et al., 2017). The BARLEYMAP pipeline was used to assign the single nucleotide polymorphisms (SNP) Array markers on the genetic map, POPSEQ_2017, and physical map, MorexV3 reference genome (Supplementary Fig. S1; Supplementary Table S1) (Cantalapiedra et al., 2015; Mascher et al., 2017; Mascher et al., 2021). The donor introgression sizes for both genetic and physical positions were calculated based on the midpoint distance between a background (GP) SNP and the flanking distal SNP of the donor introgression (Igri) (Supplementary Table S2).

GP-rapid was generated by further backcrossing the selected GP-fast line (BC3F5) to GP, followed by 4 generations of selfing with marker-assisted selection (MAS) (BC4F5). One plant from the BC4F5 population was backcrossed one additional time to GP, followed by one generation of MAS-assisted selfing (BC5F2). In the resulting BC5F2 generation, one homozygous BC5F2 line was further validated using RNAseq-based introgression genotyping (see below). This validated line was designated GP-rapid (Supplementary Fig. S2). MAS was conducted using Cleaved Amplified Polymorphic Sequences (CAPS) assays designed based on the Barley 50k iSelect SNP Array markers that differentiated the donor (Igri) and background (GP) genomes. CAPS markers were selected to cover the introgressed regions identified in GP-fast, with increased density on chromosome 2H flanking the *Ppd-H1* locus (Supplementary Table S3).

### Genotyping using RNAseq

Total RNA was extracted from leaf samples of the selected GP-rapid line, using Qiagen RNeasy Plant Mini Kit (Qiagen, Hilden, Germany, Cat. No. / ID: 74904), with the addition of 0.2 % (v/v) beta-mercaptoethanol. RNA quantity and quality were assessed with NanoPhotometer NP80 (IMPLEN, Germany) and on a 1 % agarose gel. Purified total RNA samples were sequenced at Novogene UK (Cambridge, UK). Libraries were prepared by poly-A tail capture, yielding 21 million Illumina PE150 reads (Supplementary Table S4). RNAseq data for GP and GP-fast were reported previously (Lan et al., 2025).

The RNAseq reads were quality-controlled with FastQC (v. 0.12.1) and their reports collated with MulitQC (v. 1.12) (Andrews, 2010; Ewels et al., 2016). All samples passed QC, and the RNAseq reads were aligned to the Golden Promise genome using STAR (v. 2.7.11a) aligner (Dobin et al., 2013; Schreiber et al., 2020). The following commands were applied: Genome indexing – “STAR - runMode genomeGenerate -genomeFastaFiles $genome -sjdbGTFfile $gtf -sjdbOverhang 149; RNAseq alignment – “STAR -outSAMtype BAM SortedByCoordinate -outFilterMultimapNmax 1 – outSAMmultNmax 1”.

The aligned reads (BAM) were curated with Samtools (v. 1.19.0) before calling SNPs with BCFtools (v. 1.19.0) (Danecek et al., 2021). The following commands were used: “samtools collate -O $BAM | samtools fixmate -m -- | samtools sort --o $sorted.BAM”, and SNPs were then called: “bcftools mpileup --redo-BAQ --min-BQ 30 --per-sample-mF -a AD,DP -d 1000 -f $genome -b $samples.list - Ou | bcftools call --multiallelic-caller --variants-only -Ob > out.sorted.bcf ; bcftools view -I ‘QUAL<20’ out.sorted.bcf > out.sorted.Q20.vcf”.

SNP calls in the introgression lines (GP-fast and GP-rapid) were compared to the allele called in the GP line used for backcrossing, SNPs matching the GP background were assigned 1 and 0 for alternative allele (Supplementary Table S5). The sizes of donor introgressions were calculated as before, using the midpoint distance between GP and alternative allele. The distal SNPs of introgressed regions were calculated based on a 5 Mbp rolling window, which needed to contain ≥10 SNPs that differed between the introgression lines and GP, and introgression windows closer than 15 Mbp were joined (Supplementary Table S6). Alternative allele SNP densities and introgressions were visualized and highlighted with karyoploteR (v. 1.30.0) in R (v. 4.4.3) (R Core Team, 2024; Gel and Serra, 2017).

### Growth conditions

Unless otherwise specified, plants were grown in 11*11*12 cm pots filled with the following standard substrate: Einheitserde ED73 (Einheitserde Werkverband e.V., Germany) was mixed with 7 % sand and 4 g L^−1^ Osmocote Exact Hi End, 3 to 4 M, 4th generation (ICL Group Ltd., UK). After sowing, seeds were stratified for 3 days at 4 °C and then transferred to climate-controlled growth chambers under the following conditions until maturity: 16 h light at 20 °C with photosynthetically active radiation (PAR) of 250 µmol m^−2^ s^−1^; 8 h dark, at 16 °C.

### Speed breeding conditions

Speed breeding was performed following a modified protocol based on Watson et al., 2018. Grains were first sown in 96-well trays with the standard substrate and stratified for 5 days at 4 °C. Seedlings were then grown under extended photoperiod conditions: 22 h light at 20 °C with photosynthetically active radiation (PAR) of 250 µmol m^−2^ s^−1^; 2 h dark, at 16 °C. After 15 days of cultivation, seedlings were transplanted to 11*11*12 cm pots filled with standard substrate and cultivated until maturity.

One immature spike from each plant was harvested 3 weeks post-awn tipping (Zadoks stage 49, Zadoks et al., 1974) and immediately dried at 34 °C in an air-forced oven for 4 days. Germination assays were then conducted by placing 20 grains on wetted filter papers in a petri dish. 6 dishes were prepared for each genotype. After stratifying at 4 °C for 4 days, the petri dishes were transferred to a dark incubator at 24 °C. Germination was assessed at 24h and 48h.

### Phenotypic analysis

Flowering time was recorded as the number of days after sowing (DAS) until awns emerged from the flag leaf sheath of the main culm, named awn tipping (Zadoks stage 49, Zadoks et al., 1974). Spike length, final floret number, and grain number were scored at maturity. Spike fertility was calculated as the ratio of grain number to the final floret number. Three spikes were measured for each plant.

### *Agrobacterium*-mediated barley transformation and genotyping

The binary vector construct employed to assess transformation efficiency was designed for mutagenizing the Pseudo-receiver domain of Ppd-H1 (Turner et al., 2005), using CRISPR/Cas9. The construct was cloned based on a multiplex genome editing system (Kumar et al., 2018), with two sgRNAs targeting the beginning of the 3^rd^ exon (spacer sequence: 5’ – CCCCGTCGAGAACGGCCACC – 3’) and 4^th^ exon (spacer sequence: 5’ – GATGTCGCACGATTCCA – 3’), respectively. This construct has not been previously published, and an annotated sequence file (Genbank format) can be found in Supplementary Dataset S1.

Barley immature embryo transformations followed a previously published protocol (Hensel et al., 2008), using *Agrobacterium* strain AGL1. Briefly, immature embryos (1.5-2 mm in diameter) were isolated and inoculated with *Agrobacterium* carrying the binary vector, then briefly washed and incubated at 21 °C in the dark in co-culture medium (CCM). After three days, embryos were transferred to callus induction medium (CIM) containing hygromycin (20 mg L^-1^) for selection and incubated at 21 °C in the dark. Embryos (calli) were moved to fresh CIM for an additional two weeks and subsequently to regeneration medium (RGM) with reduced hygromycin (10 mg L^-1^) under light conditions. Following two rounds of cultivation on RGM (2 weeks each), regenerated plantlets were transferred to rooting medium (RM) containing 20 mg L^-1^ hygromycin. Approximately two weeks later, plantlets were screened via PCR for successful T-DNA insertion (presence of Cas9). The following primers were used: 5’ - GAGCGCATGAAGAGGATCGA - 3’ and 5’ – GGACACGAGCTTGGACTTGA – 3’. Transgene-positive plantlets were further transferred to RM without hygromycin for further cultivation and photographing.

### Statistical analysis

Statistical tests were conducted using R (v. 4.4.3) (R Core Team, 2024). Student’s *t*-test was used for testing the significance between two groups (days to awn tipping in Fig. 2F), with a p-value cutoff at ≤ 0.01. For significance tests among three groups (days to awn tipping in Fig. 2B; spike fertility in Fig. 2D; spike length in Fig. 2E; transformation efficiency in Fig. 3C), one-way ANOVA (function *aov*) and subsequent Tukey’s HSD (function *TukeyHSD* from library *multcompView*, v. 0.1.10) were used, with a p-value cutoff at ≤ 0.05 or ≤ 0.01. Replicate numbers are indicated in the corresponding figure legends. Plots were generated using ggplot2 (v. 3.5.2) (Wickham, 2016).

**Fig. 1.**
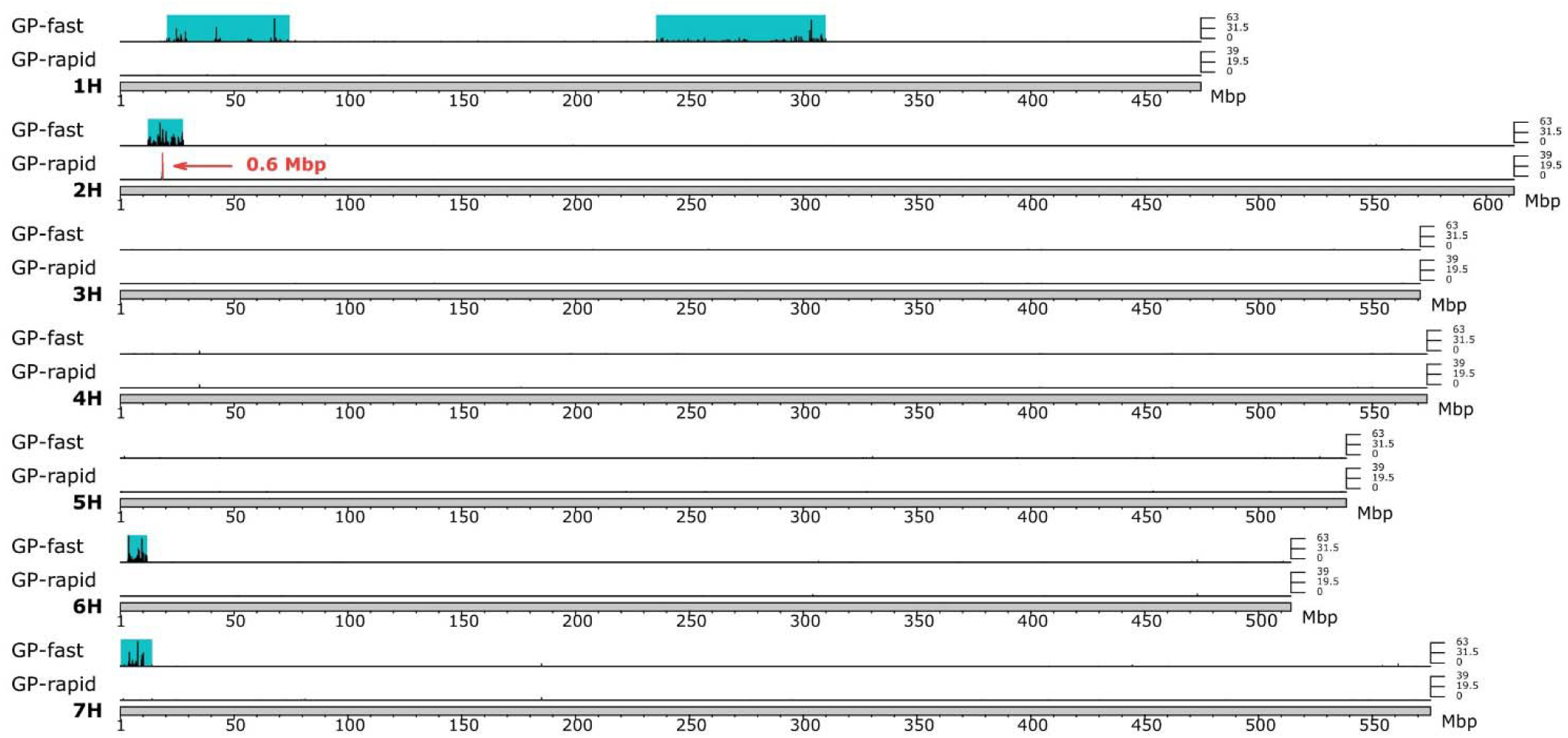
Genotyping by RNA-sequencing of Igri introgressions in GP-fast and GP-rapid. SNP density of SNPs differing GP from GP-fast (top) and GP-rapid (bottom) on the seven chromosomes of the GP genome reference, axis indicate SNP density in 10 kbp windows. Introgression regions are highlighted with rectangular boxes, GP-fast (cyan) and GP-rapid (red, location of the only introgression surrounding the *Ppd-H1* locus).

**Fig. 2.**
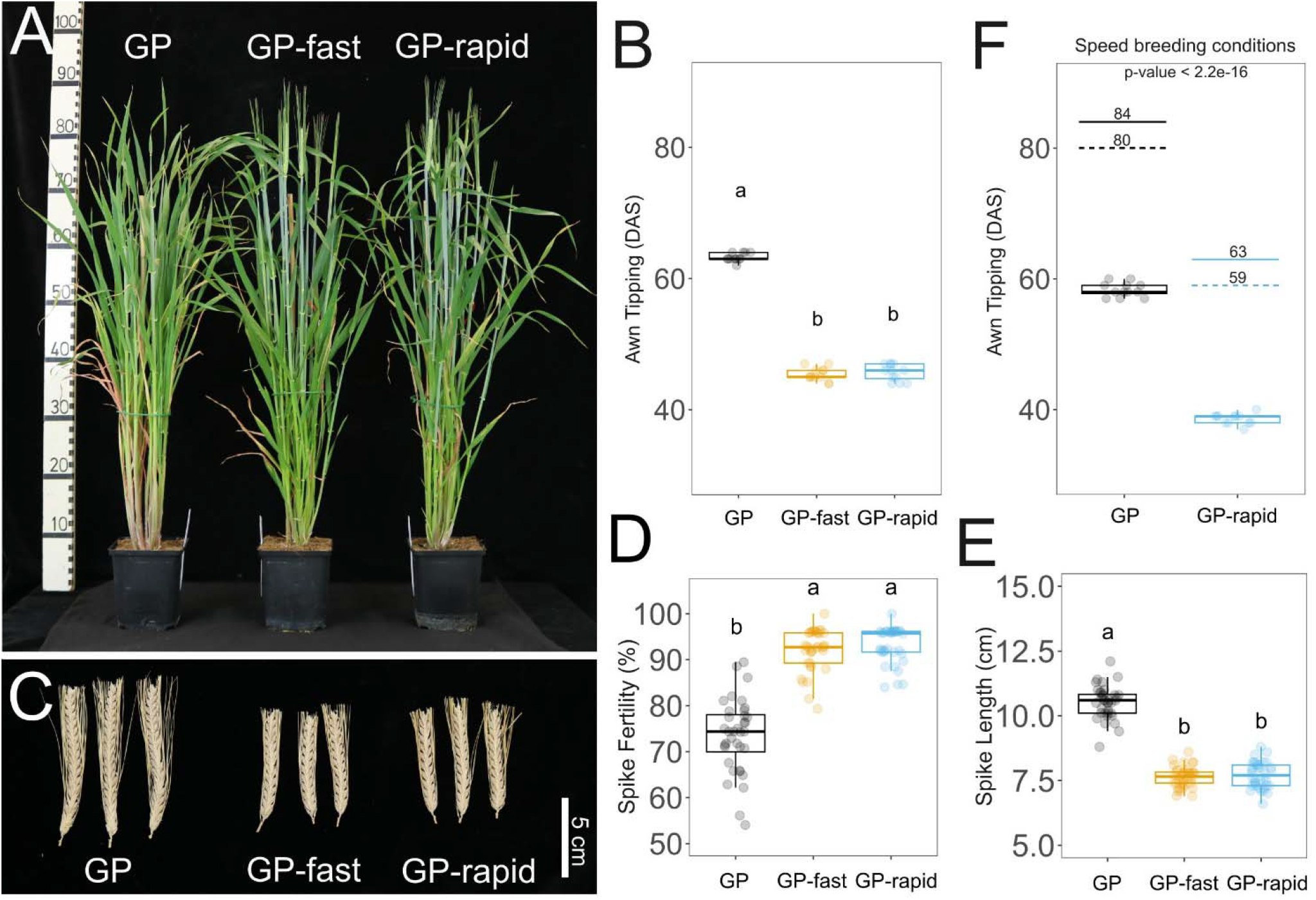
Phenotypical comparisons across GP, GP-fast, and GP-rapid under long-day (16 h light / 8 h dark) conditions (A-E) and Speed breeding (22 h light / 2 h dark) conditions (F). (A) Representative plants 61 days after sowing (DAS). Fully emerged spikes can be observed for GP-fast and GP-rapid, whereas awns are not yet visible for GP. (B) Number of days between sowing and awn tipping, n = 12. (C) Spike morphology at maturity, scale bar = 5 cm. (D) Spike fertility (Nr. Grains / Nr. Florets in percentage), and (E) spike length, n = 36. (F) Awn tipping and growing cycle comparison between GP and GP-rapid under Speed breeding conditions. Dashed lines indicate when immature spikes were harvested (three weeks post-awn tipping), and the solid lines represent when viable grains were obtained (dried at 34 °C for 4 days), n = 12. For (B), (D), and (E), the data were analyzed by a one-way ANOVA and Tukey’s honestly significant difference (HSD). Error bars represent standard deviation, p-values < 0.01. For (F), Student’s *t*-test was employed.

**Fig. 3.**
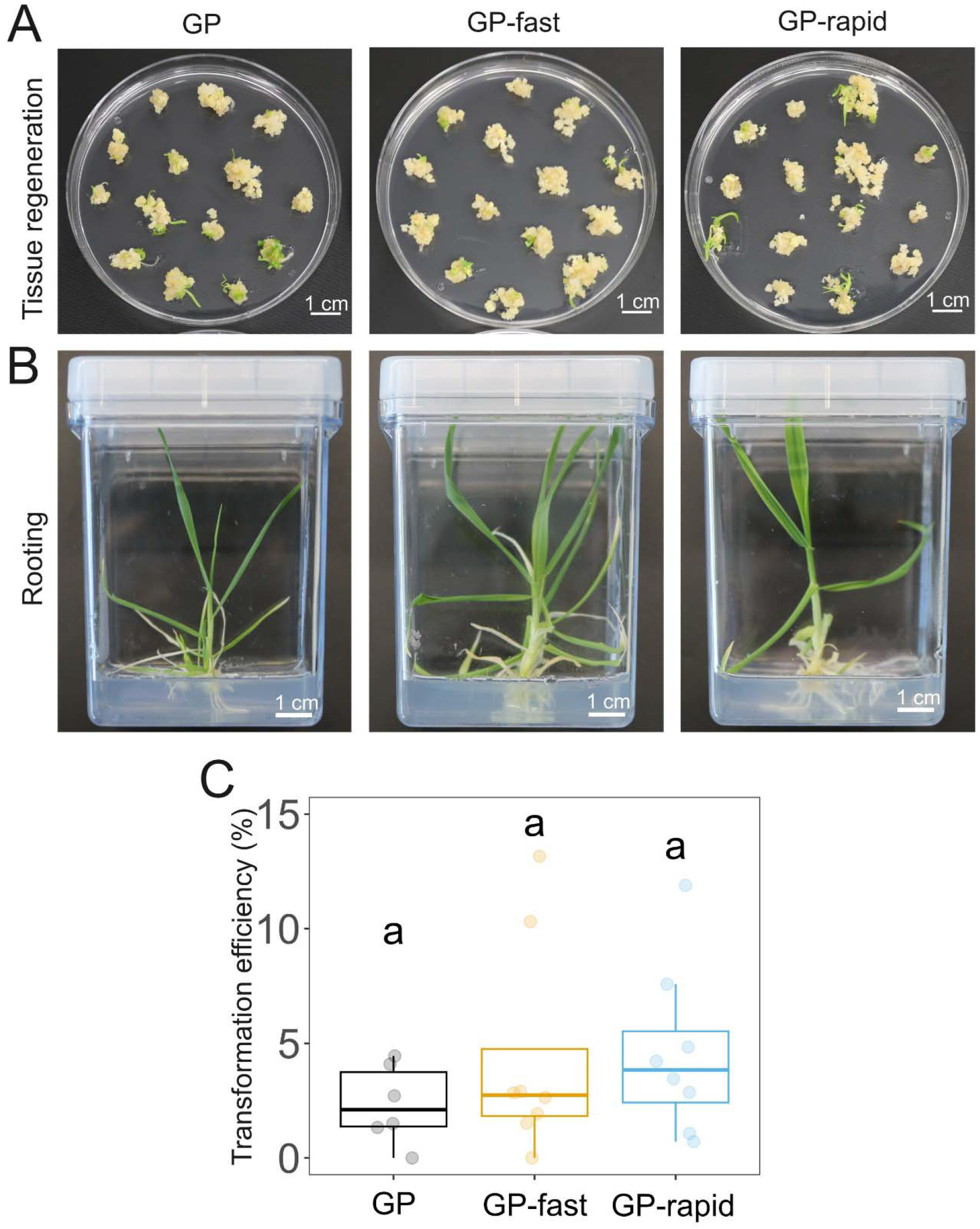
Regeneration and transformation amenability in GP, GP-fast, and GP-rapid. (A) Calli regeneration 14 days after transferring to regeneration medium (RGM) containing hygromycin, scale bars = 1 cm. (B) Representative transgene-positive plants 8 days after transferring to rooting medium (RM) without hygromycin, scale bars = 1 cm. (C) Transformation efficiency comparison across the three genotypes. Data points represent individual transformations, n = 6-8. In each transformation, the number of explants varies from 37 to 175. A detailed summary of transformation results can be found in Table 3. The data were analyzed by a one-way ANOVA and Tukey’s honestly significant difference (HSD). No significant difference was detected (p-values > 0.05). Error bars represent standard deviation.

## Results

### GP-rapid retains one introgression of approximately 0.6 Mbp flanking the *Ppd-H1* allele

Genotyping of GP-fast with the Barley 50k iSelect SNP array revealed introgressions on chromosome (Chr) 1H (41.15-48.64 cM), 2H (flanking the *Ppd-H1* locus, 12.18-41.65 cM), 6H (5.84-24.58 cM), and 7H (1.53-21.52 cM) from the Igri genome (Supplementary Fig. S1; Supplementary Table S1, S2). To reduce the identified introgressions, we next conducted two additional rounds of backcrossing to GP guided by marker-assisted selection (MAS) using Cleaved Amplified Polymorphic Sequences (CAPS) markers (Supplementary Fig. S2; Supplementary Table S3). From the BC5F2 generation, a single plant was selected – hereafter referred to as GP-rapid – that only contained a single homozygous introgression surrounding the *Ppd-H1* locus on Chr 2H. RNA sequencing confirmed that the selected GP-rapid line contained only a single introgression on Chr 2H flanking the *Ppd-H1* locus, which was reduced from 16 Mbp to approximately 0.6 Mbp (Chr 2H: 18217415-18825119) (Fig. 1; Supplementary Table S6). There were a total of 26 genes and two Long Terminal Repeats (LTRs) located within the introgressed interval. Their names, genomic positions, and putative gene functions are detailed in Table 1. Collectively, via MAS backcrossing, we generated a near-isogenic line GP-rapid, with an introgression surrounding *Ppd-H1* of 0.6 Mbp, including only 26 genes.

**Table 1.** List of genes within the retaining introgression in GP-rapid.

| Gene ID | Gene location | Putative gene function |
| --- | --- | --- |
| Horvu_GOLDEN_2H01G071200 | 18216628 - 18218107 | Pectate lyase |
| Horvu_GOLDEN_2H01G071300 | 18222613 - 18223779 | Ta11-like non-LTR retrotransposon |
| Horvu_GOLDEN_2H01G071400 | 18225883 - 18228960 | LPS-induced tumor necrosis factor alpha factor |
| Horvu_GOLDEN_2H01G071500 | 18239054 - 18241586 | Actin cross-linking protein |
| Horvu_GOLDEN_2H01G071600 | 18285599 - 18287394 | Transcription factor |
| Horvu_GOLDEN_2H01G071700 | 18293972 - 18296679 | Protein kinase-like protein |
| Horvu_GOLDEN_2H01G071800 | 18346036 - 18346605 | RING/U-box superfamily protein |
| **Horvu_GOLDEN_2H01G071900** | **18351304 - 18354525** | **Pseudo-response regulator (Ppd-H1)** |
| Horvu_GOLDEN_2H01G072000 | 18366492 - 18367622 | Strictosidine synthase |
| Horvu_GOLDEN_2H01G072100 | 18394728 - 18399897 | Sporulation protein RMD1 |
| Horvu_GOLDEN_2H01G072200 | 18401162 - 18403746 | Ascorbate peroxidase |
| Horvu_GOLDEN_2H01G072300 | 18404602 - 18407365 | Arogenate dehydratase |
| Horvu_GOLDEN_2H01G072400 | 18444402 - 18446931 | Myb/SANT-like DNA-binding domain protein |
| Horvu_GOLDEN_2H01G072500 | 18452826 - 18453248 | Tapetum determinant 1 |
| Horvu_GOLDEN_2H01G072600 | 18477588 - 18480133 | UvrABC system protein C |
| Horvu_GOLDEN_2H01G072700 | 18482260 - 18484041 | Serine/threonine-protein kinase ATM |
| Horvu_GOLDEN_2H01G072800 | 18500675 - 18502613 | Glycosyltransferases |
| Horvu_GOLDEN_2H01G072900 | 18504006 - 18507914 | Glycosyltransferases |
| Horvu_GOLDEN_2H01G073000 | 18567774 - 18569554 | Elongation factor 1-alpha |
| Horvu_GOLDEN_2H01G073100 | 18633447 - 18635137 | Retrotransposon protein, putative, unclassified |
| Horvu_GOLDEN_2H01G073200 | 18641793 - 18642503 | Leucine-rich repeat receptor-like protein kinase family protein |
| Horvu_GOLDEN_2H01G073300 | 18675353 - 18677359 | Elongation factor 1-alpha |
| Horvu_GOLDEN_2H01G073400 | 18679940 - 18684042 | Yellow stripe-like transporter 12 |
| Horvu_GOLDEN_2H01G073500 | 18692529 - 18693696 | Retrovirus-related Pol polyprotein from transposon TNT 1-94 |
| Horvu_GOLDEN_2H01G073600 | 18738300 - 18741443 | MLO-like protein |
| Horvu_GOLDEN_2H01G073700 | 18763915 - 18766983 | MLO-like protein |
| Horvu_GOLDEN_2H01G073800 | 18816650 - 18817055 | GRF zinc finger family protein |
| Horvu_GOLDEN_2H01G073900 | 18817600 - 18820898 | Phosphoinositide phospholipase C |

### GP-rapid displays early flowering and improved spike fertility

We compared the developmental traits of GP-rapid with GP and GP-fast. All plants were grown under LD conditions with 16 h light/8 h dark. GP-rapid displayed an early flowering phenotype compared to GP (Fig. 2A). Awn tipping (Zadoks stage 49, Zadoks et al., 1974) in GP-rapid occurred on average 45.7 days after sowing, 18 days earlier than GP, but was not significantly different from GP-fast, which flowered on average 45.4 days after sowing (Fig. 2B). Additionally, we also observed that GP-rapid and GP-fast had comparable and shorter spikes (7.6±0.4 cm and 7.7±0.5 cm, respectively) than GP (10.5±0.6 cm) (Fig. 2C, E). Despite a reduction in spike length, GP-rapid displayed a higher spike fertility (93.2 %±3.9) compared to GP (73.4 %±8.0), similar to GP-fast (92.1 %±4.6) (Fig. 2C, D). These observations are consistent with previous reports, indicating that the wild-type *Ppd-H1* allele is associated with higher spike fertility under different conditions (Digel et al., 2015; Ejaz and von Korff, 2017; Gol et al., 2021; Lan et al., 2025)

Overall, these findings show that GP-rapid is early-flowering and displays higher spike fertility as compared to GP, providing further evidence that the *Ppd-H1* allele is primarily responsible for the trait differences described for GP-fast.

### GP-rapid exhibits a 25 % shorter growth cycle under speed breeding conditions

The advent of speed-breeding strategies has further revolutionized crop development timelines by significantly reducing generation times in various crops, including barley (Watson et al., 2018). These methods combine extended photoperiods (e.g., 22 h Light/2 h Dark) and the use of pre-harvest immature spikes, enabling up to six generations per year, compared to the conventional three generations under standard growth conditions (Watson et al., 2018). Two genes, *Ppd-H1* and *ELF3*, have been identified as key contributors to accelerated development under speed-breeding conditions (Rossi et al., 2024). Given that GP-rapid carries the wild-type *Ppd-H1* allele, its flowering time and generation time could likely be further reduced through speed-breeding protocols.

To test this, we further compared flowering time and seed-to-seed cycle time between GP-rapid and GP under speed breeding conditions. Our results showed that under extended photoperiods (22 h light/2 h dark), awn tipping in GP-rapid took place 38.6 days after sowing (Fig. 2F), about 7 days shorter than when grown under 16 h Light/8 h Dark (45.7 days, Fig. 2B). GP, however, reached the same developmental stage about 20 days later than GP-rapid under speed breeding condition (58.3 days, Fig. 2F). Both genotypes progressed to early dough stage (Zadoks stage 83, Zadoks et al., 1974) three weeks post-awn tipping, at which point immature spikes were harvested (Fig. 2F, dashed lines), dried at 34 °C for 4 days, and subjected to germination tests (Fig. 2F, solid lines). Germination rates exceeded 80 % in both genotypes within 48 hours (Table 2), confirming the viability of harvested seeds.

**Table 2.**
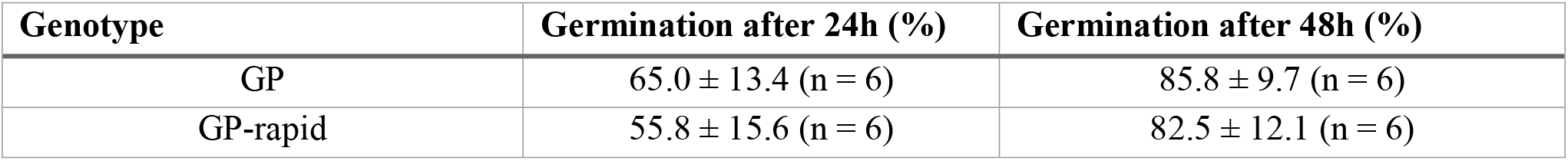
Germination test of grains harvested from the speed breeding trial.

| Genotype | Germination after 24h (%) | Germination after 48h (%) |
| --- | --- | --- |
| GP | 65.0 ± 13.4 (n = 6) | 85.8 ± 9.7 (n = 6) |
| GP-rapid | 55.8 ± 15.6 (n = 6) | 82.5 ± 12.1 (n = 6) |

**Table 3.**
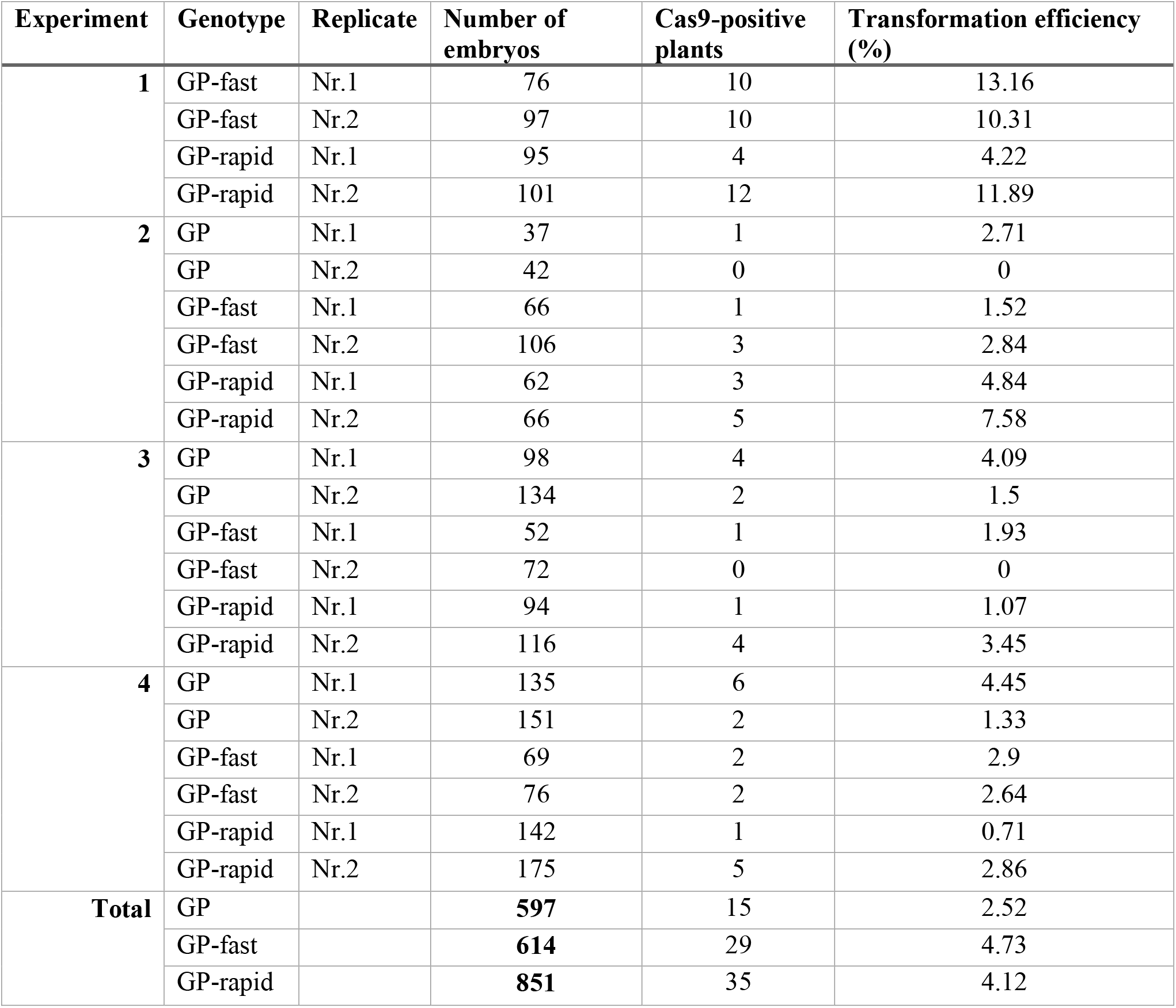
Transformation efficiencies using the Ppd-H1 CRISPR construct.

| **Experiment** | **Genotype** | **Replicate** | **Number of embryos** | **Cas9-positive plants** | **Transformation efficiency (%)** |
| --- | --- | --- | --- | --- | --- |
| **1** | GP-fast | Nr.1 | 76 | 10 | 13.16 |
|  | GP-fast | Nr.2 | 97 | 10 | 10.31 |
|  | GP-rapid | Nr.1 | 95 | 4 | 4.22 |
|  | GP-rapid | Nr.2 | 101 | 12 | 11.89 |
| **2** | GP | Nr.1 | 37 | 1 | 2.71 |
|  | GP | Nr.2 | 42 | 0 | 0 |
|  | GP-fast | Nr.1 | 66 | 1 | 1.52 |
|  | GP-fast | Nr.2 | 106 | 3 | 2.84 |
|  | GP-rapid | Nr.1 | 62 | 3 | 4.84 |
|  | GP-rapid | Nr.2 | 66 | 5 | 7.58 |
| **3** | GP | Nr.1 | 98 | 4 | 4.09 |
|  | GP | Nr.2 | 134 | 2 | 1.5 |
|  | GP-fast | Nr.1 | 52 | 1 | 1.93 |
|  | GP-fast | Nr.2 | 72 | 0 | 0 |
|  | GP-rapid | Nr.1 | 94 | 1 | 1.07 |
|  | GP-rapid | Nr.2 | 116 | 4 | 3.45 |
| **4** | GP | Nr.1 | 135 | 6 | 4.45 |
|  | GP | Nr.2 | 151 | 2 | 1.33 |
|  | GP-fast | Nr.1 | 69 | 2 | 2.9 |
|  | GP-fast | Nr.2 | 76 | 2 | 2.64 |
|  | GP-rapid | Nr.1 | 142 | 1 | 0.71 |
|  | GP-rapid | Nr.2 | 175 | 5 | 2.86 |
| **Total** | GP |  | **597** | 15 | 2.52 |
|  | GP-fast |  | **614** | 29 | 4.73 |
|  | GP-rapid |  | **851** | 35 | 4.12 |

In summary, the implementation of speed breeding enabled a complete seed-to-seed cycle in only 63 days for GP-rapid, representing a 25 % reduction compared to GP. This accelerated cycling allows for up to six generations per year and positions GP-rapid as a highly efficient genotype for fast-forwarding genetic studies.

### GP-rapid is amenable to transformation

We next investigated whether GP-rapid maintains amenability to *Agrobacterium*-mediated immature embryo transformation as GP. Previous studies identified Transformation Amenability (TFA) loci contributing to GP’s high transformation amenability (Hisano and Sato, 2016; Hisano et al., 2017; Orman-Ligeza et al., 2020). Since GP-rapid still retains a segment from the Igri genome on Chr 2H (Fig. 1), which is in proximity to one of the TFA loci (TFA5, Hisano et al., 2016), it was critical to evaluate the transformation efficiency of GP-rapid alongside GP to compare their transformation amenability. To do so, we chose a CRISPR construct targeting the Pseudo-receiver domain of Ppd-H1 and performed *Agrobacterium*-mediated transformation using immature embryos from GP and GP-rapid, as well as GP-fast. Four independent transformations were conducted. For each genotype, the total number of embryos was between 597 and 851 (Table 3).

Efficient regeneration from explants is a critical requirement for successful transformation. In our trials, all three genotypes exhibited rapid plantlet regeneration from calli within two weeks on regeneration medium containing hygromycin (selective agent) (Fig. 3A). Furthermore, transgenic plantlets developed healthy shoots and roots on the rooting medium (Fig. 3B). Genotyping of T0 plants revealed no statistically significant differences in transformation efficiencies among the three genotypes (Fig. 3C). Notably, when data from all transformations were pooled, we observed slightly higher overall transformation efficiency in GP-rapid (4.12 %) and GP-fast (4.73 %) compared to GP (2.52 %) (Table 3). Prior reports have suggested that transformation success is influenced by the physiological condition of donor plants, with stress-free plants producing higher-quality explants (Harwood, 2014; Hayta et al., 2021). Considering that the wild-type *Ppd-H1* allele increases robustness against stresses during reproductive development (Gol et al., 2021; Lan et al., 2025), the modestly increased transformation efficiency observed in GP-rapid and GP-fast may reflect improved explant quality compared to GP. In conclusion, GP-rapid remains highly amenable to immature embryo transformation.

## Discussion

Recent advancements in CRISPR/Cas-mediated genome editing technologies have facilitated precise and efficient targeted mutagenesis in numerous crop species, including barley (Gasparis et al., 2018; Lawrenson et al., 2021; Lawrenson et al., 2024). Golden Promise (GP), known for its high transformation amenability and well-characterized genome, has become the reference genotype for genetic transformation (Murray et al., 2004; Hensel et al., 2008; Harwood, 2012; Schreiber et al., 2020). In this study, we generated a fast-cycling GP isogenic homozygous line, named GP-rapid (Fig. 1). This genotype showed 25 % shorter seed-to-seed cycle time compared to GP under speed breeding conditions, enabling up to six generations per year (Fig. 2F). Importantly, the Igri introgression was significantly reduced relative to GP-fast, simplifying downstream applications such as guide RNA design for genome editing.

Beyond shortening the life cycle, the wild-type *Ppd-H1* allele in barley enhances resilience and stabilizes development and yield under abiotic stress conditions (Wiegmann et al., 2019; Gol et al., 2021; Lan et al., 2025). Furthermore, lines carrying the wild-type *Ppd*-H1 allele maintained transcriptomic stability, in contrast to spring barley genotypes with the natural *ppd-H1* mutation, which activated a strong transcriptional stress response under high-temperature conditions (Lan et al., 2025). Therefore, GP-rapid may be particularly advantageous for transformation if donor or transgenic plants are grown in environments with suboptimal or variable growth conditions.

In conclusion, GP-rapid is a versatile and efficient genetic system for barley research. It combines a fast generation cycle, high transformation efficiency, and improved stress tolerance. Its adoption may substantially accelerate functional genomic studies and genome editing applications in barley.

## Supplementary data

The following supplementary data are available at JXB online.

Table S1. Igri introgressions in the selected single GP-fast line using the Barley 50k iSelect SNP array.

Table S2. Igri introgression sizes in the selected single GP-fast line based on the Barley 50k iSelect SNP array.

Table S3. List of CAPS Markers used for marker-assisted genotype selection. Table S4. RNAseq libraries used for genotyping of introgressions.

Table S5. Igri introgression analysis using RNAseq-based genotyping in the selected single GP-fast line and GP-rapid based on RNAseq-based genotyping.

Table S6. Igri introgression sizes in the selected single GP-fast line and GP-rapid based on RNAseq-based genotyping.

Fig. S1. Barley 50k iSelect SNP array genotyping of the selected GP-fast line used for further backcrossing with GP.

Fig. S2. Graphic overview of GP-rapid generation.

Dataset S1. An annotated sequence file of the plasmid used for testing transformation efficiency.

## Acknowledgments

We are grateful to Gesa Helmsorig for her support in designing the CAPS markers and to Nina Döring for her assistance with plant cultivation.

## Author contributions

GB, EBH, SL, and MvK conceived and designed the research. RS performed the crossings; EBH and TR carried out marker-assisted genotyping; EBH analyzed the Barley 50k iSelect SNP array and RNAseq-based genotyping data; SL conducted phenotypic comparisons; AW and GB designed and cloned the plasmid for transformation efficiency testing. SL and GB performed the transformation experiments; SL, EBH, and MvK drafted and revised the manuscript, with input from all authors.

## Data availability

The RNA-Seq data used for genotyping are available in DataPLANT at [URL], and can be accessed with [deposition number XXX]. All other data supporting the findings of this work are available within the article and its online Supplementary data.

---

**Fig. S1.**
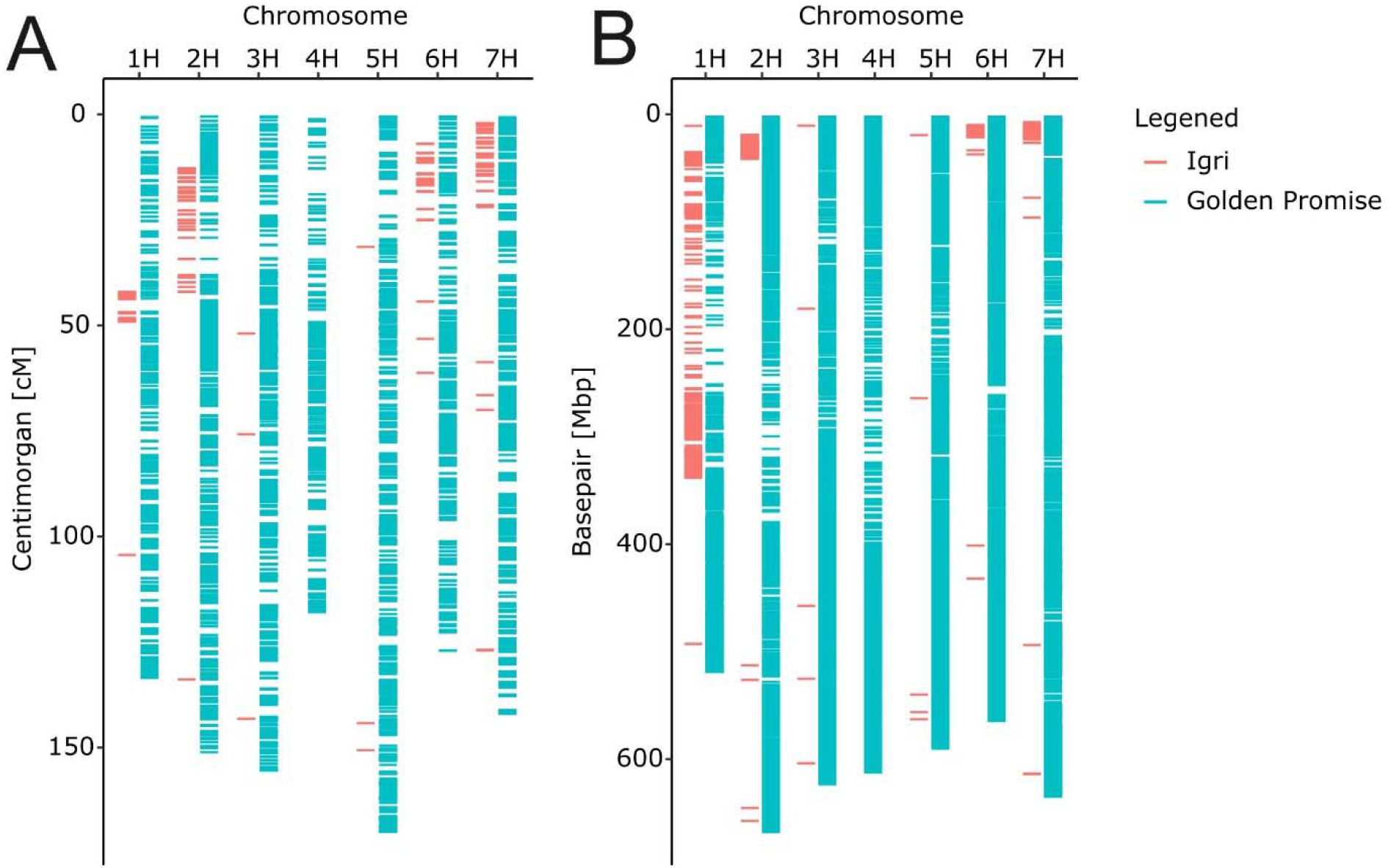
Barley 50k iSelect SNP array genotyping of the selected GP-fast line (GP-fast_9) used for further backcrossing with GP. Markers are indicated for Igri (red, foreign material) and GP background in (blue). (A) Genetic map with marker location from POPSEQ_2017 and (B) physical map with marker physical location obtained from Morex V3 reference genome (Cantalapiedra et al., 2015).

**Fig. S2.**
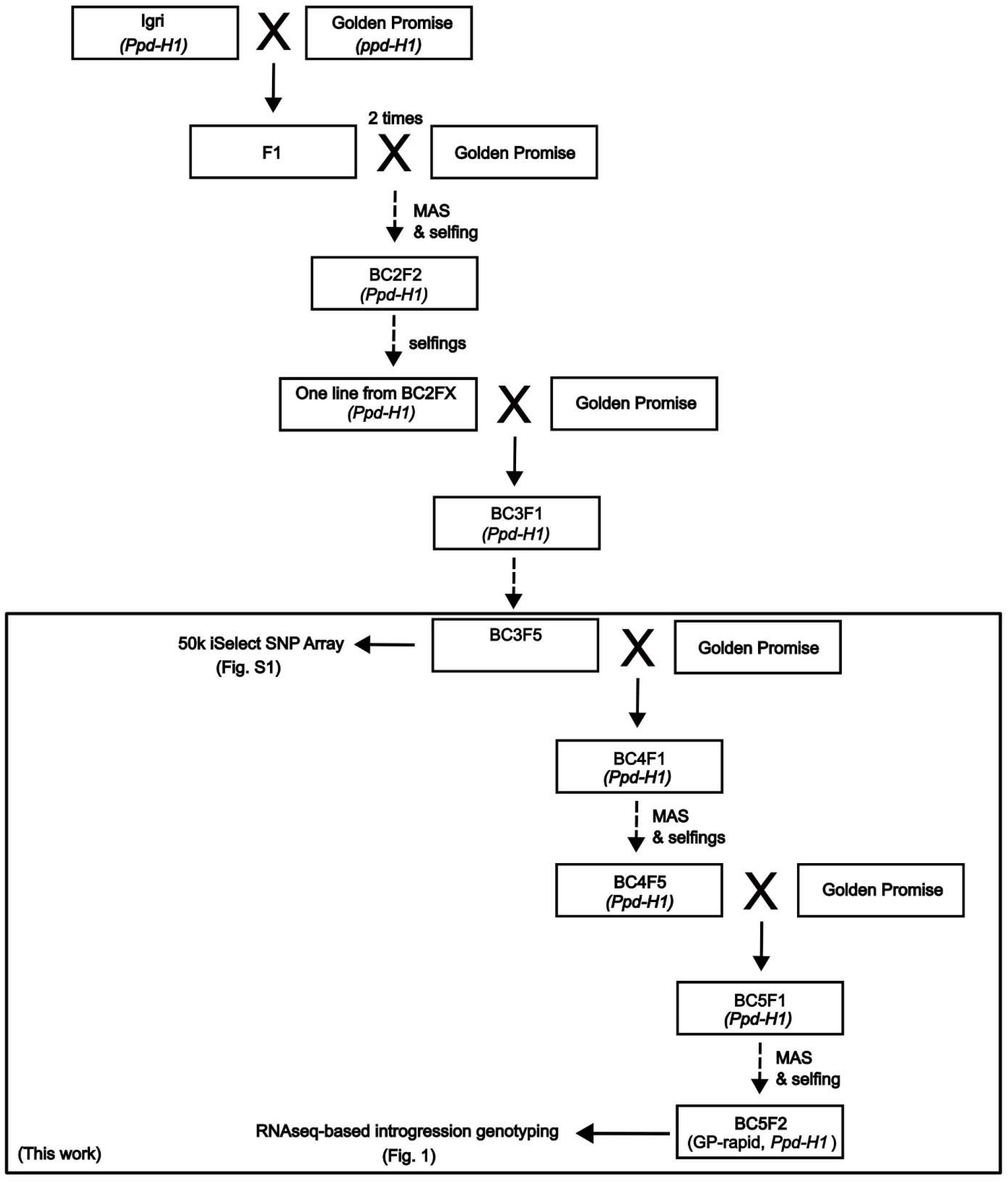
Graphic overview of GP-rapid generation.

## References

Amanda D, Frey FP, Neumann U, et al. 2022. Auxin boosts energy generation pathways to fuel pollen maturation in barley. Current Biology 32, 1798-1811.e8.

Andrews S. 2010. FastQC: A Quality Control tool for High Throughput Sequence Data. Babraham Bioinformatics, Cambridge, UK.

Bayer MM, Rapazote-Flores P, Ganal M, et al. 2017. Development and Evaluation of a Barley 50k iSelect SNP Array. Frontiers in Plant Science 8.

Campoli C, Drosse B, Searle I, Coupland G, von Korff M. 2012. Functional characterisation of HvCO1, the barley ( Hordeum vulgare ) flowering time ortholog of CONSTANS. The Plant Journal 69, 868–880.

Cantalapiedra CP, Boudiar R, Casas AM, Igartua E, Contreras-Moreira B. 2015. BARLEYMAP: physical and genetic mapping of nucleotide sequences and annotation of surrounding loci in barley. Molecular Breeding 35, 13.

Cockram J, Donini P, Chiapparino E, Laurie DA, Stamati K, Taylor SA, O’Sullivan DM. 2007. Haplotype analysis of vernalization loci in European barley germplasm reveals novel VRN-H1 alleles and a predominant winter VRN-H1/VRN-H2 multi-locus haplotype. Theoretical and Applied Genetics 115, 993–1001.

Danecek P, Pollard MO, Liddle J, et al. 2021. Twelve years of SAMtools and BCFtools. GigaScience 10.

Digel B, Pankin A, von Korff M. 2015. Global Transcriptome Profiling of Developing Leaf and Shoot Apices Reveals Distinct Genetic and Environmental Control of Floral Transition and Inflorescence Development in Barley. The Plant Cell 27, 2318–2334.

Dobin A, Davis CA, Schlesinger F, Drenkow J, Zaleski C, Jha S, Batut P, Chaisson M, Gingeras TR. 2013. STAR: ultrafast universal RNA-seq aligner. Bioinformatics 29, 15–21.

Ejaz M, Von Korff M. 2017. The Genetic Control of Reproductive Development under High Ambient Temperature. Plant Physiology 173, 294–306.

Ewels P, Magnusson M, Lundin S, Käller M. 2016. MultiQC: summarize analysis results for multiple tools and samples in a single report. Bioinformatics 32, 3047–3048.

Forster BP. 2001. Mutation genetics of salt tolerance in barley: An assessment of Golden Promise and other semi-dwarf mutants. Euphytica 120, 317–328.

Gasparis S, Kała M, Przyborowski M, Łyżnik LA, Orczyk W, Nadolska-Orczyk A. 2018. A simple and efficient CRISPR/Cas9 platform for induction of single and multiple, heritable mutations in barley (Hordeum vulgare L.). Plant Methods 14, 111.

Gel B, Serra E. 2017. karyoploteR: an R/Bioconductor package to plot customizable genomes displaying arbitrary data. (J Hancock, Ed.). Bioinformatics 33, 3088–3090.

Gol L, Haraldsson EB, von Korff M. 2021. Ppd-H1 integrates drought stress signals to control spike development and flowering time in barley. (M Jones, Ed.). Journal of Experimental Botany 72, 122–136.

Han Y, Broughton S, Liu L, Zhang X-Q, Zeng J, He X, Li C. 2021. Highly efficient and genotype-independent barley gene editing based on anther culture. Plant Communications 2, 100082.

Harwood WA. 2012. Advances and remaining challenges in the transformation of barley and wheat. Journal of Experimental Botany 63, 1791–1798.

Harwood WA. 2014. A Protocol for High-Throughput Agrobacterium-Mediated Barley Transformation. In: Henry RJ, Furtado A, eds. Cereal Genomics. Totowa, NJ: Humana Press, 251– 260.

Harwood WA (Ed.). 2019. Barley: Methods and Protocols. New York, NY: Springer New York.

Hayta S, Smedley MA, Clarke M, Forner M, Harwood WA. 2021. An Efficient Agrobacterium LJMediated Transformation Protocol for Hexaploid and Tetraploid Wheat. Current Protocols 1, e58.

Helmsorig G, Walla A, Rütjes T, Buchmann G, Schüller R, Hensel G, von Korff M. 2024. early maturity 7 promotes early flowering by controlling the light input into the circadian clock in barley. Plant Physiology 194, 849–866.

Hensel G, Valkov V, Middlefell-Williams J, Kumlehn J. 2008. Efficient generation of transgenic barley: The way forward to modulate plant–microbe interactions. Journal of Plant Physiology 165, 71–82.

Hisano H, Meints B, Moscou MJ, Cistue L, Echávarri B, Sato K, Hayes PM. 2017. Selection of transformation-efficient barley genotypes based on TFA (transformation amenability) haplotype and higher resolution mapping of the TFA loci. Plant Cell Reports 36, 611–620.

Hisano H, Sato K. 2016. Genomic regions responsible for amenability to Agrobacterium-mediated transformation in barley. Scientific Reports 6, 37505.

Jayakodi M, Padmarasu S, Haberer G, et al. 2020. The barley pan-genome reveals the hidden legacy of mutation breeding. Nature 588, 284–289.

Kumar N, Galli M, Ordon J, Stuttmann J, Kogel K, Imani J. 2018. Further analysis of barley MORC 1 using a highly efficient RNA LJguided Cas9 geneLJediting system. Plant Biotechnology Journal 16, 1892–1903.

Lan T, Walla A, Çolpan Karışan KE, et al. 2025. PHOTOPERIOD 1 enhances stress resistance and energy metabolism to promote spike fertility in barley under high ambient temperatures. Plant Physiology 197, kiaf118.

Lawrenson T, Clarke M, Kirby R, Forner M, Steuernagel B, Brown JKM, Harwood W. 2024. An optimised CRISPR Cas9 and Cas12a mutagenesis toolkit for Barley and Wheat. Plant Methods 20, 123.

Lawrenson T, Hinchliffe A, Clarke M, Morgan Y, Harwood W. 2021. In-planta Gene Targeting in Barley Using Cas9 With and Without Geminiviral Replicons. Frontiers in Genome Editing 3, 663380.

Marthe C, Kumlehn J, Hensel G. 2015. Barley (Hordeum vulgare L.) Transformation Using Immature Embryos. In: Wang K, ed. Agrobacterium Protocols: Volume 1. New York, NY: Springer, 71–83.

Mascher M, Gundlach H, Himmelbach A, et al. 2017. A chromosome conformation capture ordered sequence of the barley genome. Nature 544, 427–433.

Mascher M, Wicker T, Jenkins J, et al. 2021. Long-read sequence assembly: a technical evaluation in barley. The Plant Cell 33, 1888–1906.

Mulki MA, Bi X, von Korff M. 2018. FLOWERING LOCUS T3 Controls Spikelet Initiation But Not Floral Development. Plant Physiology 178, 1170–1186.

Müller Y, Patwari P, Stöcker T, et al. 2023. Isolation and characterization of the gene HvFAR1 encoding ACYLLJCOA reductase from the cer□za.227 mutant of barley ( Hordeum vulgare ) and analysis of the cuticular barrier functions. New Phytologist 239, 1903–1918.

Murray F, Brettell R, Matthews P, Bishop D, Jacobsen J. 2004. Comparison of Agrobacterium - mediated transformation of four barley cultivars using the GFP and GUS reporter genes. Plant Cell Reports 22, 397–402.

Orman-Ligeza B, Harwood W, Hedley PE, Hinchcliffe A, Macaulay M, Uauy C, Trafford K. 2020. TRA1: A Locus Responsible for Controlling Agrobacterium-Mediated Transformability in Barley. Frontiers in Plant Science 11, 355.

Pieper R, Tomé F, Pankin A, von Korff M. 2021. FLOWERING LOCUS T4 delays flowering and decreases floret fertility in barley. (Z Wilson, Ed.). Journal of Experimental Botany 72, 107–121.

R Core Team. 2024. R: A language and environment for statistical computing. R Foundation for Statistical Computing, Vienna, Austria. URL https://www.R-project.org/.

Rossi N, Powell W, Mackay IJ, Hickey L, Maurer A, Pillen K, Halliday K, Sharma R. 2024. Investigating the genetic control of plant development in spring barley under speed breeding conditions. Theoretical and Applied Genetics 137, 115.

Saisho D, Takeda K. 2011. Barley: Emergence as a New Research Material of Crop Science. Plant and Cell Physiology 52, 724–727.

Schreiber M, Mascher M, Wright J, Padmarasu S, Himmelbach A, Heavens D, Milne L, Clavijo BJ, Stein N, Waugh R. 2020. A Genome Assembly of the Barley ‘Transformation Reference’ Cultivar Golden Promise. G3 Genes|Genomes|Genetics 10, 1823–1827.

Tamilselvan-Nattar-Amutha S, Hiekel S, Hartmann F, Lorenz J, Dabhi RV, Dreissig S, Hensel G, Kumlehn J, Heckmann S. 2023. Barley stripe mosaic virus-mediated somatic and heritable gene editing in barley (Hordeum vulgare L.). Frontiers in Plant Science 14, 1201446.

Turner A, Beales J, Faure S, Dunford RP, Laurie DA. 2005. The Pseudo-Response Regulator Ppd-H1 Provides Adaptation to Photoperiod in Barley. Science 310, 1031–1034.

Vardanega I, Maika JE, Demesa-Arevalo E, et al. 2025. CLAVATA signalling shapes barley inflorescence by controlling activity and determinacy of shoot meristem and rachilla. Nature Communications 16, 3937.

Wanke A, Van Boerdonk S, Mahdi LK, et al. 2023. A GH81-type β-glucan-binding protein enhances colonization by mutualistic fungi in barley. Current Biology 33, 5071-5084.e7.

Watson A, Ghosh S, Williams MJ, et al. 2018. Speed breeding is a powerful tool to accelerate crop research and breeding. Nature Plants 4, 23–29.

Wendt T, Hansson M, Dockter C, Waugh R, Holme I, Druka A, Braumann I, Preuß A, Thomas W. 2016. HvDep1 Is a Positive Regulator of Culm Elongation and Grain Size in Barley and Impacts Yield in an Environment-Dependent Manner. PLOS ONE 11, e0168924.

Wickham H. 2016. ggplot2: Elegant Graphics for Data Analysis. Springer-Verlag New York. ISBN 978-3-319-24277-4, https://ggplot2.tidyverse.org.

Wiegmann M, Maurer A, Pham A, et al. 2019. Barley yield formation under abiotic stress depends on the interplay between flowering time genes and environmental cues. Scientific Reports 9, 6397.

Zadoks JC, Chang TT, Konzak CF. 1974. A decimal code for the growth stages of cereals. Weed Research 14, 415–421.

